# GeneSetCluster 2.0: a comprehensive toolset for summarizing and integrating gene-sets analysis

**DOI:** 10.1101/2024.12.18.629178

**Authors:** Asier Ortega-Legarreta, Alberto Maillo, Daniel Mouzo, Ana Rosa López-Pérez, Lara Kular, Majid Pahlevan Kakhki, Maja Jagodic, Jesper Tegner, Vincenzo Lagani, Ewoud Ewing, David Gomez-Cabrero

## Abstract

**Background:** Gene-Set Analysis (GSA) is commonly used to analyze high-throughput experiments. However, GSA cannot readily disentangle clusters or pathways due to redundancies in upstream knowledge bases, which hinders comprehensive exploration and interpretation of biological findings. To address this challenge, we developed GeneSetCluster, an R package designed to summarize and integrate GSA results. Over time, we and users as well identified limitations in the original version, such as difficulties in managing redundancies across multiple gene-sets, large computational times, and its lack of accessibility for users without programming expertise.

**Results:** We present GeneSetCluster 2.0, a comprehensive upgrade that delivers methodological, computational, interpretative, and user-experience enhancements. Methodologically, GeneSetCluster 2.0 introduces a novel approach to address duplicated gene-sets and implements a seriation-based clustering algorithm that reorders results, aiding pattern identification. Computationally, the package is optimized for parallel processing, significantly reducing execution time. GeneSetCluster 2.0 enhances cluster annotations by associating clusters with relevant tissues and biological processes to improve biological interpretation, particularly for human and mouse data. To broaden accessibility, we have developed a user-friendly web application enabling non-programmers to use it. This version also ensures seamless integration between the R package, catering to users with programming expertise, and the web application for broader audiences. We evaluated the updates in a single-cell RNA public dataset.

**Conclusion:** GeneSetCluster 2.0 offers substantial improvements over its predecessor. Furthermore, by bridging the gap between bioinformaticians and clinicians in multidisciplinary teams, GeneSetCluster 2.0 facilitates collaborative research. The R package and web application, along with detailed installation and usage guides, are available on GitHub (https://github.com/TranslationalBioinformaticsUnit/GeneSetCluster2.0), and the web application can be accessed at https://translationalbio.shinyapps.io/genesetcluster/.

## 1. Background

High-throughput technologies are fundamental for understanding biological systems by enabling the profiling of thousands to millions of features (e.g., genes) on a genome-wide scale. The final stages of their associated bioinformatic analysis aim to identify the system’s most relevant features. However, interpreting these features in a biological context can be overwhelming (1). To address these challenges, methods have been developed to identify functionally related groups of features, offering biologists higher-order, interpretable summaries of their experiments (2).

Gene-Set Analysis (GSA) has emerged as the standard for functional bioinformatic analysis in gene expression studies. Methodologically, two main approaches dominate GSA: Over-Representation Analysis (ORA (3)) and Gene-Set Enrichment Analysis (GSEA (4)). Briefly, ORA determines whether the proportion of relevant genes in a gene-set exceeds the expected by chance. At the same time, GSEA ranks all genes based on their association with a trait and tests whether genes within a particular set cluster toward the top of the ranking, reflecting their importance. Beyond gene expression, tools such as GREAT (5) extend GSA to other genomic features, including DNA methylation and chromatin accessibility, by first linking genomic ranges to genes. Equally critical to the methodologies are the gene-sets, curated from existing scientific literature or derived from molecular experiments. Prominent resources include the Gene Ontology (GO) project (6), Reactome (7), the Kyoto Encyclopedia of Genes and Genomes (KEGG) pathway database (8), and the Molecular Signatures Database (MSigDB) (9).

Despite the utility of GSA, and regardless of the methodology used, interpreting its results remains challenging. First, the focus shifts from interpreting individual genes to interpreting gene-sets or pathways, but GSA often identifies thousands of overlapping processes, complicating the interpretation. This redundancy derives from gene-set overlap, where highly related pathways are repeatedly significant, resulting in top-ranked processes that reflect the same underlying signal. Second, researchers frequently need to “analyze multiple contrasts within a single study” (e.g., screening various drugs against controls or performing *knock outs*) producing extensive lists of overlapping gene-sets, either across contrasts or from different databases, or “multiple gene-sets derived from the same contrast but from several gene-set databases”, or both. Several approaches have been developed to improve GSA interpretability, which we will denote by GSA interpretation tools (GSAit). “Slim Ontologies” reduce redundancy by collapsing databases into discrete categories (9–11) while other methods incorporate the graph structure of databases like GO into statistical frameworks (12–14). However, both frameworks are database-specific and not widely generalizable. A more versatile, data-driven framework (15–20) defines distances between gene-sets (e.g., based on shared genes or semantic similarity), clusters them using these distances, and interprets the clusters through text mining or representative gene-sets.

We initially developed GeneSetCluster 1.0 (21), a GSAit tool to address these challenges. It measured distances between gene-sets using relative risk (RR) and applied hierarchical clustering to identify clusters of gene-sets. However, despite its successful application (22,23) the tool had limitations, including non-interpretable clusters caused by identical or “outlier” gene-sets, and an inability to refine clustering despite identifying associated challenges. Furthermore, GeneSetCluster 1.0 was available only as an R package, restricting its use to bioinformatics experts with programming skills.

GeneSetCluster 2.0 introduces several significant enhancements to overcome the limitations of its predecessor. Methodologically, it incorporates a novel approach to address duplicated gene-sets and utilizes a seriation-based clustering algorithm to reorder results, facilitating the identification of meaningful patterns. Computationally, the tool is optimized for parallel processing, which significantly reduces execution times and enhances efficiency. To improve biological interpretation, particularly for human and mouse data, GeneSetCluster 2.0 enriches cluster annotations by associating clusters with relevant tissues and biological processes, enabling more comprehensive insights. Additionally, the tool broadens accessibility through developing a user-friendly web application, allowing non-programmers to leverage its functionality while maintaining the R package for users with programming expertise. The web-application version also ensures seamless integration between the R package.

Together, these improvements make GeneSetCluster 2.0 a robust and versatile solution for Gene-Set Interpretation Analysis, enabling a diverse user base and facilitating better integration of bioinformatics into multidisciplinary research workflows.

## 2. Implementation

In this section we first briefly describe version 1.0 of GeneSetCluster. Then we separately illustrate the new functionalities implemented in version 2.0 for the gene-set cluster identification and interpretation.

### 2.1 GeneSetCluster 1.0

GeneSetCluster 1.0 (21) is an R package designed to address a critical challenge in gene-set analysis (GSA): interpreting results that often encompass hundreds to thousands of potentially overlapping gene-sets. Furthermore, a common issue in GSA is that many gene-sets represent similar biological processes but are labeled differently, making it challenging to identify overarching themes.

GeneSetCluster 1.0 addresses this challenge by grouping gene-sets based on shared genes, using RR as the distance metric, and employing k-means or hierarchical clustering methods. To determine the optimal number of clusters, the package uses silhouette analysis (24) and the elbow method (25). Notably, this approach assigns all input gene-sets to a cluster, which could be considered a limiting factor. To facilitate interpretation, GeneSetCluster 1.0 enables the gene-set cluster visualization in three visualization schemes: as a network, as a dendrogram, or as a heatmap.

In summary, GeneSetCluster 1.0 (v1.0) enables the integration of results from different GSA tools and experimental conditions, offering a unified framework for exploring multiple GSA results simultaneously. For extended details, we refer to the original publication (21).

### 2.2 GeneSetCluster 1.0 limitations in the clustering analysis

Several limitations have been identified over the past years of user experience with version 1.0. The first limitation was related to clustering analysis. While sub-clusters could be visually identified after a clustering analysis, the tool did not allow for re-clustering these sub-groups to achieve greater granularity.

Second, multiple GSA results often identified the same gene-sets (e.g., identical Gene Ontology IDs), despite slight variations in the subsets of genes associated with each result. In GeneSetCluster 1.0, these duplicated gene-sets were treated independently—a methodology we refer to as the “Raw Gene-Sets” approach. This occasionally introduced bias into the clustering process, leading, for example, to the largest cluster being composed of these repeated gene-sets. Third, the clustering methods used—k-means and hierarchical clustering—forced each gene-set into a cluster. This constraint limited the biological interpretability of the resulting clusters.

To address these limitations, we implemented three modules - sub-clustering analysis, merging duplicated gene-sets, and seriation analysis.

#### Sub-clustering analysis

GeneSetCluster 2.0 implements *BreakUpCluster*, which enables selecting a gene-set cluster and identifying sub-clusters within it (“*breaking it*” into smaller sub-clusters) (Fig. 2). This targeted refinement addresses the issue of the challenging interpretation of large clusters. By allowing researchers to focus on specific clusters of interest, *BreakUpCluster* provides a detailed exploration of finer gene-set relationships while preserving the overall clustering framework.

#### Merging duplicated gene-sets

Multiple GSA results often identify the same gene-sets (e.g., identical Gene Ontology IDs), even though the subsets of genes associated with each GSA result may vary slightly. In GeneSetCluster 1.0, these duplicated gene-sets were treated independently, an approach we refer to as the “Raw Gene-Sets” methodology. This approach occasionally introduced bias into the clustering process, sometimes resulting in the largest cluster being dominated by these repeated gene-sets, which were distinct from other clusters in the analysis.

To address this limitation, GeneSetCluster 2.0 introduces a new approach called “Unique Gene-Sets.” This method detects repeated gene-sets with identical ID labels and merges them into a single, unified entry that contains the union of all genes associated with these sets. For example, GO:0007612 (a biological process related to “learning”) might be identified in one GSA analysis due to the genes *Pak6*, *Reln*, and *Adcy3*, and in another due to *Reln*, *Adcy3*, and *Eif2ak4*. The “Unique Gene-Sets” methodology merges these results, counting GO:0007612 only once and consolidating the associated genes into a single list: *Pak6*, *Reln*, *Adcy3*, and *Eif2ak4*. Consequently, each gene-set is treated as a unique entity during the clustering process, eliminating the bias caused by duplications (Fig. 1a).

**Fig 1.**
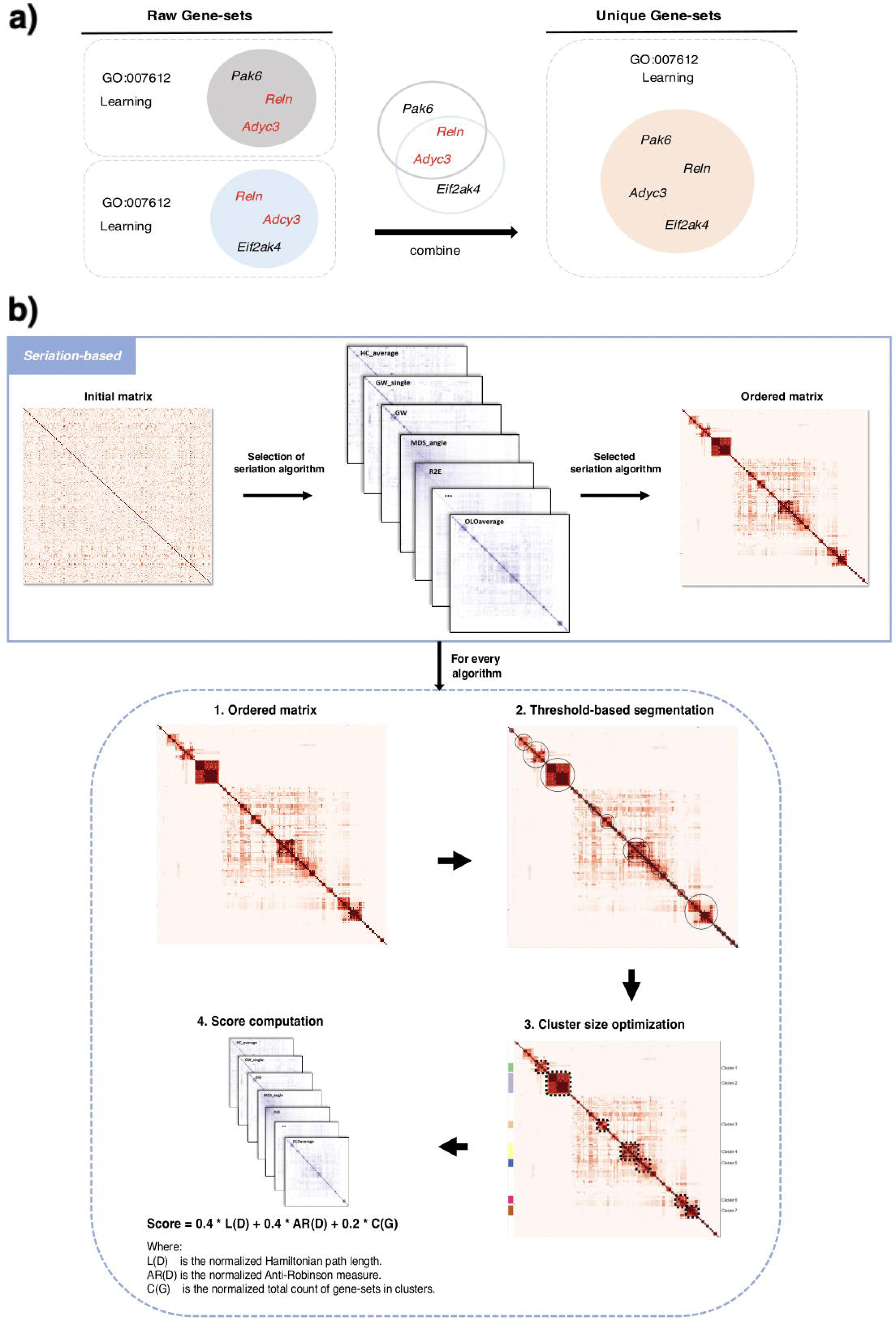
New implementations of GeneSetCluster 2.0. (A) Methods for handling duplicate gene-sets. Left side illustrates “*Raw Gene-sets*” method, considering all input gene-sets as separate entities. Right side depicts the new “*Unique Gene-sets*” approach, which merges duplicate gene-sets by combining their genes. (B) Visualization of the “*Seriation-based*” method.

This refined approach simplifies the analysis, resulting in more precise and interpretable clusters. It also facilitates more consistent comparisons across studies and enhances the understanding of the biological significance across different research contexts.

#### Seriation-based clustering approach

GeneSetCluster 2.0 enhances its predecessor by offering a new “*seriation-based*” clustering approach. Briefly, seriation methods aim to reorder data, typically rows and columns of a similarity or distance matrix, to uncover patterns that might not be apparent otherwise. These methods offer improvements over k-means or hierarchical clustering in certain contexts by focusing on the relative order of elements rather than assigning them into discrete clusters or nested hierarchies. For consistency, the k-means and hierarchical methods remain available in the updated version.

Generally, seriation-based (25) clustering identifies coherent groups by arranging gene-sets in a linear sequence based according to their pairwise similarities. Unlike k-means or hierarchical clustering, which strictly divides gene-sets into separate groups, seriation emphasizes uncovering patterns within the data (26). Specifically, GeneSetCluster 2.0 tests 32 seriation algorithms (Table S1) from the *seriation* R package (27) to automatically find the optimal algorithm (Fig. 1b). Briefly, each seriation algorithm undergoes a four-step evaluation process:

1. <u>Initial Seriation</u>: The distance matrix, which represents the pairwise similarities between gene-sets, is reordered by the algorithm, optimizing the placement of similar gene-sets next to each other.
2. <u>Threshold-Based Segmentation</u>: Potential clusters are identified within the reordered matrix by applying a predefined similarity threshold (default set at 0.6). This process groups adjacent gene-sets with similarity scores above this threshold. The similarity threshold can be manually adjusted by the users.
3. <u>Cluster Size Optimi</u>z<u>ation</u>: Clusters are refined to meet a specified size requirement (minimum size from 4 to 10 gene-sets per cluster) ensuring each cluster contains a meaningful number of gene-sets. This requirement prevents excessive fragmentation and over-large clusters that may overlook biologically relevant relationships. Users can also modify the minimum size threshold manually.
4. <u>Score computation</u>: The output of each algorithm is scored based on three weighted criteria: Hamiltonian path length (minimizing distances between consecutive gene-sets) (28), anti-Robinson form criterion ensuring dissimilarities increase when moving away from the diagonal (29), and a total number of clustered gene-sets (maximizing the number of gene-sets successfully grouped), weighted at 40%, 40%, and 20%, respectively (see Supplementary material for details).

Finally, the seriation algorithm with the highest score from step 4 is automatically selected as the optimal solution. However, users can override this automatic selection by specifying their preferred algorithm. Similarly, the optimal minimum number of gene-sets required to define a cluster is automatically determined but can be modified by users.

The “*Seriation-based*” method provides several key advantages: 1) it effectively isolates outlier gene-sets that share low similarity with others, preventing them from introducing noise into the clustering results; 2) the optimal seriation algorithm is automatically selected based on objective scoring criteria, minimizing user intervention; and 3) it can detect gradual transitions or hierarchical structures that might be overlooked by traditional clustering methods (see *Results section*). This new method provides a more detailed and comprehensive understanding of the gene-sets relationships.

In summary, GeneSetCluster 2.0 incorporates both “*Classic*” (k-means, hierarchical) and “*Seriation-based”* clustering techniques, providing a robust toolkit for exploring and understanding complex gene-set relationships. The “*Classic*” methods can discover well-defined patterns, while the “*Seriation-based*” approach reveals subtle transitions or hierarchical structures.

### 2.3 Functional annotations

In v1.0, interpretation of gene-set clusters were delegated to the user (e.g. investigating user supplied gene subset) or using plugins to WebgestaltR (30) and StringDB (31). In v2.0, we aim to allow the interpretation of data-driven within the tool. A key concept used during annotation is *cluster-associated genes*, which correspond to the union of the genes in all gene-sets in a specific cluster.

#### Automatic Gene-Set Cluster Annotations

Within v2.0 each gene-set cluster can be annotated through two approaches. In the first approach, v2.0 applies ORA to the cluster-associated *genes,* using the *enrichGO* function from the *clusterProfiler* R package for this analysis (32). In the second approach, GeneSetCluster 2.0 conducts a semantic enrichment analysis for Gene Ontology (GO) using the *simplifyGO* function from the *SimplifyEnrichment* package (33). Internally, *simplifyGO* extracts biological themes by computing how closely related GO terms are in the GO hierarchy. This process clarifies the biological context of each cluster by emphasizing functional themes within the GO terms. This meta-analysis helps researchers to determine whether particular pathways or processes are enriched in the cluster-associated genes.

#### Tissue enrichment

GeneSetCluster 2.0 incorporates a tissue enrichment analysis (limited to human data) that identifies possible associations between gene-set clusters with human tissues. v2.0 conducts a GSEA (4) per gene-set cluster where the gene-set is the *cluster-associated genes*, and the ranking per tissue is based on the gene expression available in the GTEx database (34), which contains median expression levels across 54 human tissues. As a result, we obtain a ranking of relevant tissues per gene-set cluster. For computational efficiency, users can access the tissue expression database via the API integrated into GeneSetCluster 2.0 or by downloading directly from the repository for local use.

### 2.4 Computational enhancements

GeneSetCluster 1.0 faced limitations in computational scalability, particularly when processing large numbers of gene-sets. GeneSetCluster 2.0 incorporates a parallelization scheme to address this issue to enhance computational efficiency. This improvement is especially evident in the CombineGeneSets step (Fig. 2), where distances between gene-sets are calculated. Parallel processing techniques were implemented using the R packages doParallel (35) and foreach (36). These tools enable the computational workload to be distributed efficiently across multiple processors or threads. As a result, GeneSetCluster 2.0 achieves faster execution times and enhanced overall performance, making it well-suited for handling large-scale datasets.

**Fig 2.**
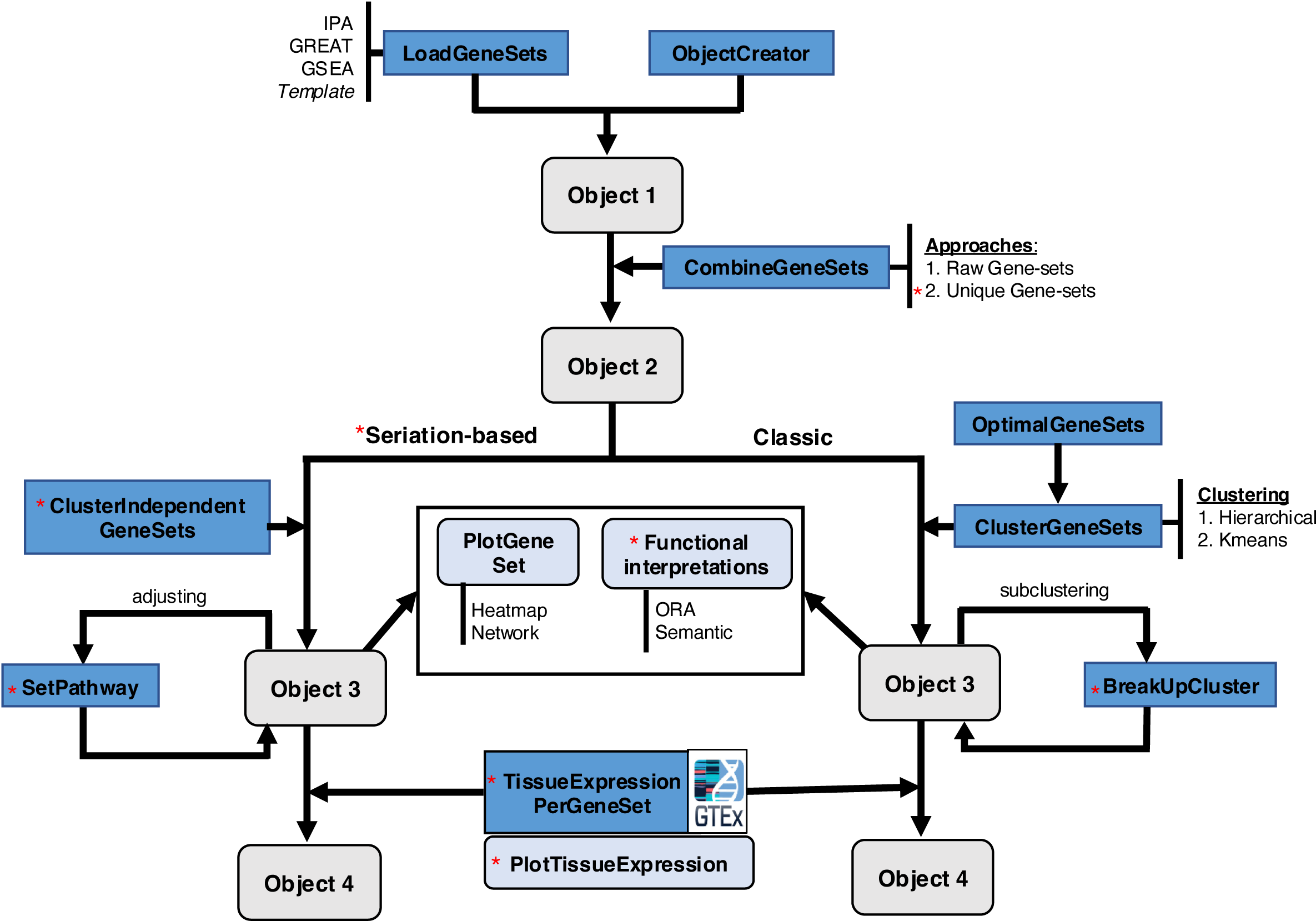
Workflow of GeneSetCluster 2.0. The illustration outlines all the features, with new implementations indicated by red asterisks. After uploading the GSA results, users can choose between two methods for handling duplicated gene sets: “*Raw Gene-sets*” and “*Unique Gene-sets*” during the *CombineGeneSet* step. Subsequently, the new “*Seriation-based*” clustering approach can be applied. Within this clustering method, the function *ClusterIndependentGeneSets* calculates the groups based on an optimal number, which can be manually adjusted using *SetPathway*. On the other hand, with the “*Classic*” method, clusters can now be subdivided into smaller clusters using *BreakUpCluster*. Finally, the outputs of both clustering methods can be annotated with additional functional interpretations (ORA and *Wordcloud*) or through tissue enrichment analysis using the *TissueExpressionPerGeneSet* function and its corresponding plot, *PlotTissueExpression*.

## 3. Web Application

To enable experienced non-programming users, we developed a web-based Shiny application. Overall, we consider the Web Application to be part of the v2.0 improvements described in Section 2. The Shiny application, developed with R 4.3, is hosted on a Shiny server at the URL https://translationalbio.shinyapps.io/genesetcluster/. It is fully compatible with all operating systems and web browsers. Upon accessing the application, users are prompted to select the mandatory input parameters and upload their GSA results. Once the “*Run Analysis*” button is clicked, the results, along with corresponding plots, are generated and displayed. Users can then examine the results, add additional annotations, and perform further analyses to gain biological insights. All results and plots are available for download. Additionally, users can save their analysis by downloading an *RData* file for future use, allowing them to continue by re-uploading it in the Shiny application or moving to the R package version. This transition can also be made from the R package to the Shiny application. For a more user-friendly experience, the application can also be deployed locally from the R package using the command *run_app()* (Fig. 3).

**Fig 3.**
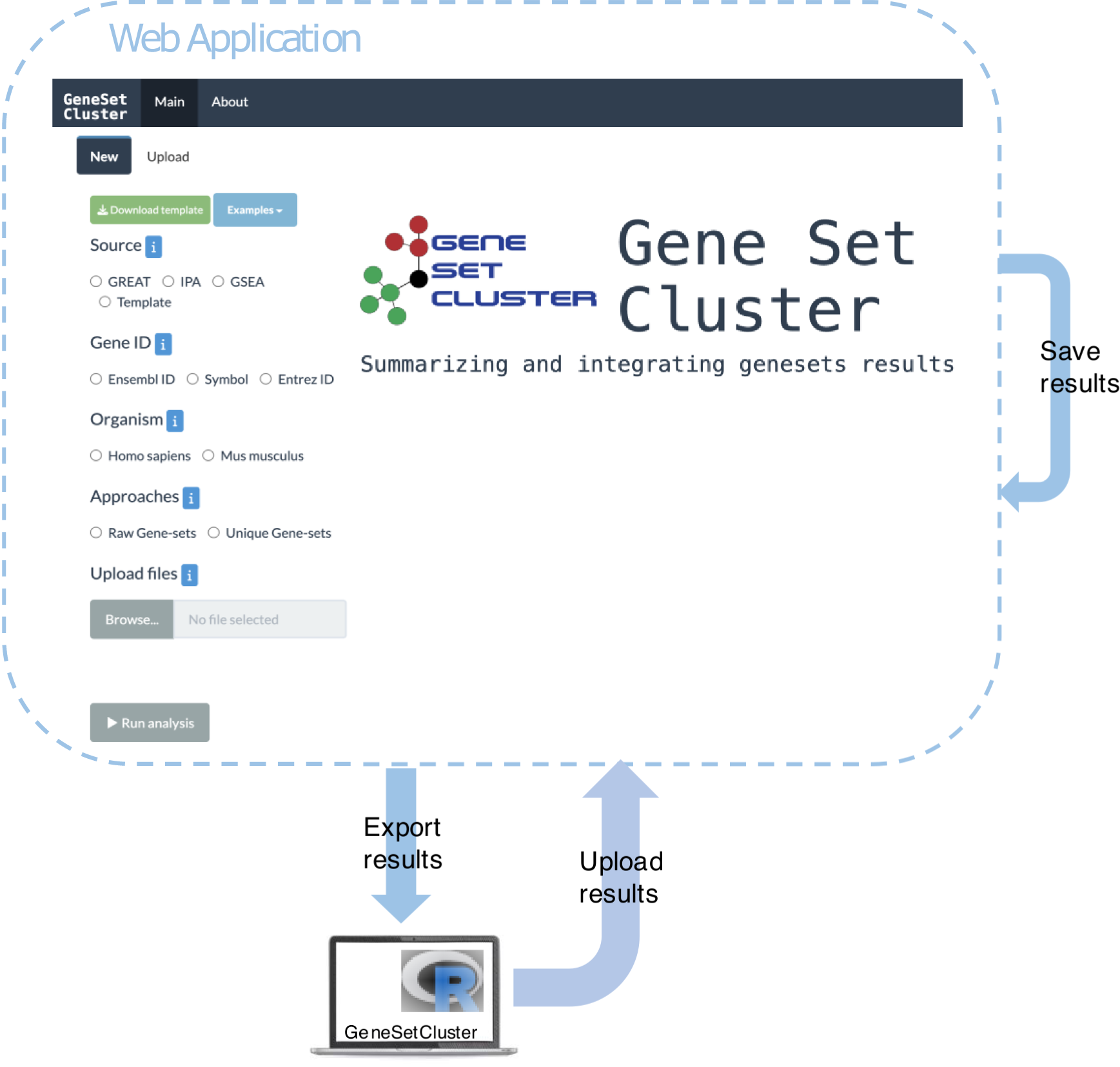
General overview of the interaction. Overview of the seamless integration between the web application and R package.

### 3.1 Graphical user interface

The user interface of the web application is divided into two main sections, as illustrated in Fig. 4. In the input section, users specify parameter values such as *Source*, *Gene ID…* Users can also start from a previously saved analysis or import an analysis performed using the R package.

**Fig 4.**
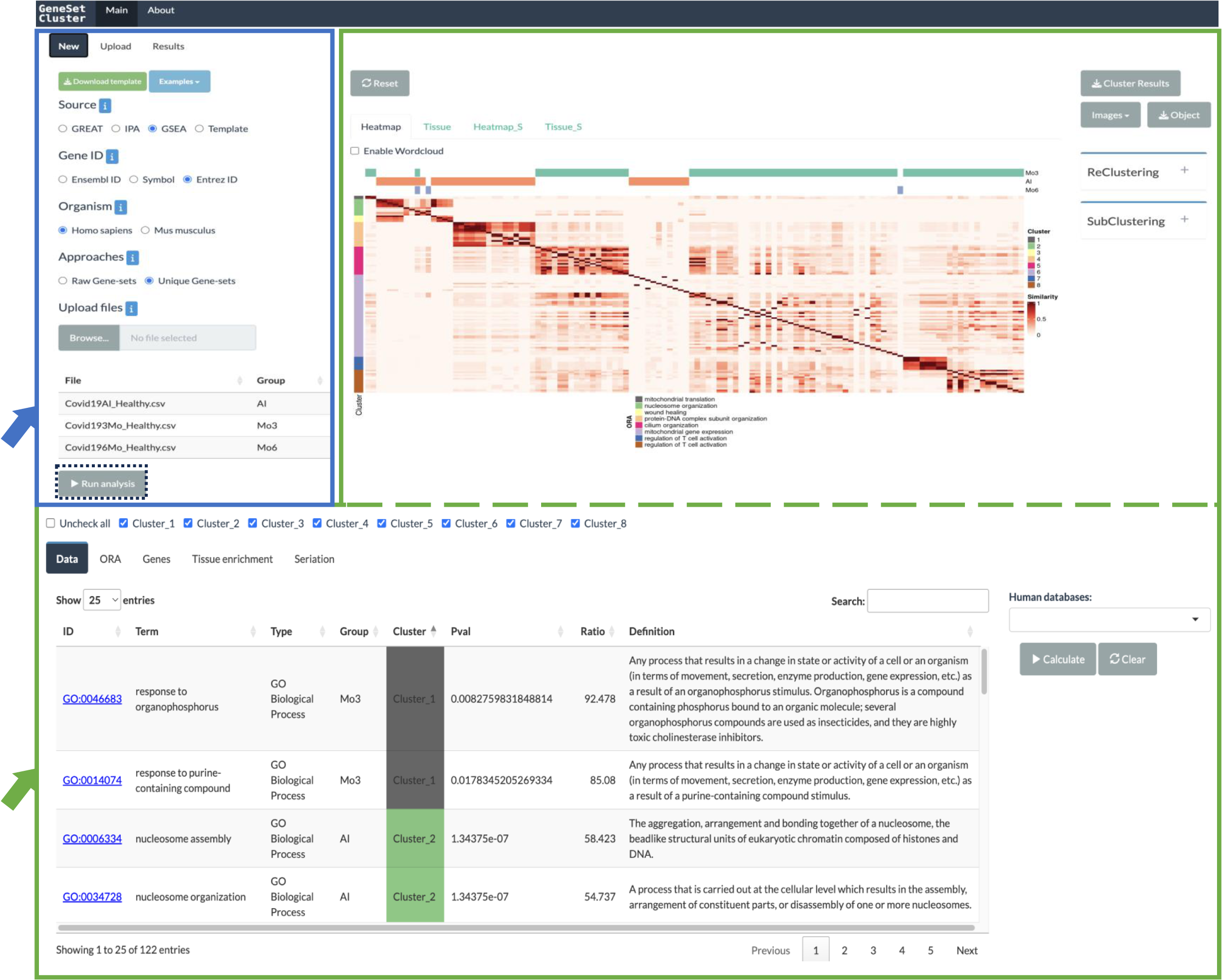
Web application interface of GeneSetCluster 2.0. The screenshot presents two main sections. The input section, highlighted in blue, allows the user to specify parameter values. After the user clicks on “*Run Analysis*” the result section will display, highlighted in green. This section displays the results in tables at the bottom and plots at the top.

The results section is further divided into two parts. At the top, users can view different plots, including heatmaps of the RR matrix (under the *Heatmap_S* tab for the “*Seriation-based*” clustering and under *Heatmap* tab for the other clustering methods), along with their tissue enrichment plots (available in the *Tissue_S* and *Tissue* tabs respectively). Below the plots, the results are displayed in tables, showing which gene-sets belong to each cluster (under the *Data* tab), ORA details (under the *ORA* tab), gene information (under the *Genes* tab), and tissue data (under the *Tissue enrichment* tab). The source code of the application can be accessed at https://github.com/TranslationalBioinformaticsUnit/GeneSetCluster2.0.

### 3.2 Analysis workflow

The application supports input files from GREAT, IPA, and GSEA in various formats, including *.csv*, *.tsv*, and *Excel*. For other tools, users can use our provided *template*, available in *Excel* format through the application, to input the necessary information for GeneSetCluster compatibility. The required fields in the template are: *ID* (gene-set identifier), *Count* (number of genes from your list found in the gene-set), *GeneRatio* (calculated as the number of genes found divided by the total genes in the gene-set), *p.adjust* (adjusted p-value), and *geneID* (list of genes in Ensembl ID, SYMBOL or ENTREZ ID). Additionally, regardless of the input source, users must specify complementary information, including the gene ID format, organism, and the preferred method for handling duplicates.

Once the data are uploaded and the user specifies their preferences, clicking the “*Run analysis*” button initiates the default analysis using the k-means method. The results, including corresponding plots, are displayed, with the input gene-sets initially classified into clusters determined by the *OptimalGeneSets* function (Fig. 2). User can later customize the number of clusters or subdivide larger clusters into smaller subclusters for more detailed exploration.

Furthermore, the users can perform the “*Seriation-based*” method (section 2.4) or apply the tissue enrichment analysis (section 2.5) to gain further insights.

We also provide interactive features that allow users to explore the results:

#### Targeted analysis

Users may want to investigate specific conditions or phenotypes in their data, focusing on a particular group of genes. This functionality allows them to assess whether their *cluster-associated genes* are enriched in these specific gene groups. Users can either import a custom gene list or select a phenotype from well-known databases for enrichment analysis. Two key databases are accessible through the application: 1) the Human Phenotype Ontologies database (HPO), which provides standardized terms for human disease-related phenotypes (37), and 2) the Mammalian Phenotype (MP) database, which offers terms for annotating mouse phenotype data (38). Overall, this feature helps uncover relevant genetic insights by emphasizing specific groups of genes within the *cluster-associated genes*.

#### Gene-set cluster functional characterization

Users may required to explore data at the gene level rather than just focusing on gene-sets. In this case, the application displays a table listing all *cluster-associated genes* and the frequency of each gene in the cluster. This allows them to filter genes based on frequency, helping them identify the most prevalent genes within each cluster, discover genes exclusive to a specific cluster, or search for a particular gene. Moreover, an ORA can be conducted on the filtered gene list to streamline the biological interpretation. Additionally, each gene is linked to external databases such as GeneCards for human data (39) and Mouse Genome Informatics for mice (40), providing further context about gene functions and implications. This feature enhances flexibility and depth in gene exploration, allowing users to focus on the most relevant genes and obtain more detailed biological insights.

Finally, all results can be downloaded directly from the application, and the plots can be in several formats, including *.jpg*, *.png*, and .*pdf*. Furthermore, users can save their analysis as an *RData* file, allowing them to quickly resume their work by re-uploading it to the application or importing it into the R package for further analysis. This functionality fosters effective interaction and collaboration among multidisciplinary teams, improves workflow efficiency, and supports ongoing research efforts.

In summary, GeneSetCluster 2.0 is accessible online and allows users to perform all its functions in a user-friendly manner. A comprehensive and informative user guide is available at the *About* tab from https://translationalbio.shinyapps.io/genesetcluster/.

## 4. Results

### 4.1 Enhancing biological interpretation

To evaluate the performance of GeneSetCluster 2.0 and compare it with GeneSetCluster 1.0, as well as a custom manual analysis, we applied both frameworks to a publicly available single-cell RNA dataset (41). The selected dataset explored the molecular basis of myelodysplastic syndromes with a deletion of the long arm of chromosome 5, del(5q), by analyzing the transcriptional and regulatory landscape of CD34+ progenitor cells using single-cell RNA-seq. Additionally, the study evaluates the impact of lenalidomide treatment on transcriptional alterations.

The authors conducted differential expression analyses across five comparisons: 1) Non-del(5q) cells of complete responders versus at diagnosis; 2) Non-del(5q) cells of partial responders versus at diagnosis; 3) Del(5q) cells of partial responders versus non-responders; 4) Non-del(5q) cells of complete responders versus healthy cells; and, 5) Non-del(5q) cells of partial responders versus healthy cells. Complete responders refer to patients who achieved complete cytogenetic response, while partial responders refer to those with partial cytogenetic response to lenalidomide treatment. From each comparison, GSEA was conducted, identifying 20 distinct gene-sets: 12 gene-sets were found in the first three comparisons, while the remaining 8 were identified in the last two comparisons. The authors manually grouped them into 7 clusters based on the biological implications: 1) Cluster 1: Related to ubiquitin processes (7 gene-sets) 2) Cluster 2: Focused on proteasome-mediated processes (2 gene-sets). 3) Cluster 3: Linked to autophagy (2 gene-sets) 4) Cluster 4: Erythropoietin signaling (2 gene-sets). 5) Cluster 5: PD-L1/PD-1 checkpoint pathway (1 gene-set) 6) Cluster 6: Phosphatidylinositol signaling system (1 gene-set). 7) Cluster 7: Mitochondrial and ribosomal translation (8 gene-sets). These original results are visualized in Figure 7a of the publication (41).

GeneSetCluster 1.0 and GeneSetCluster 2.0 were applied to these gene-sets. In the previous version, using “*Raw Gene-sets*” and k-means/hierarchical clustering methods, only two large clusters emerged: one associated with *mitochondrial translation* (8 gene-sets) and another with *proteasome-mediated processes* (12 gene-sets). In contrast, GeneSetCluster 2.0, applying “*Unique Gene-sets*” and “*Seriation-based*” methods, generated four distinct clusters: 1) Cluster 1: Mitochondrial translation (8 gene-sets). 2) Cluster 2: protein polyubiquitination (4 gene-sets) 3) Cluster 3: Proteasome-mediated processes (2 gene-sets) 4) Cluster 4: Autophagosome assembly (2 gene-sets). Three gene-sets were left not clustered, as the new method does not force gene-sets into clusters and requires a minimum of two pathways per cluster. The biological significance of each cluster was determined through an ORA of the *cluster-associated genes* against the biological process database. Fig. 5 illustrates these results. This analysis can be performed using both the R package and the web version of the tool.

**Fig 5.**
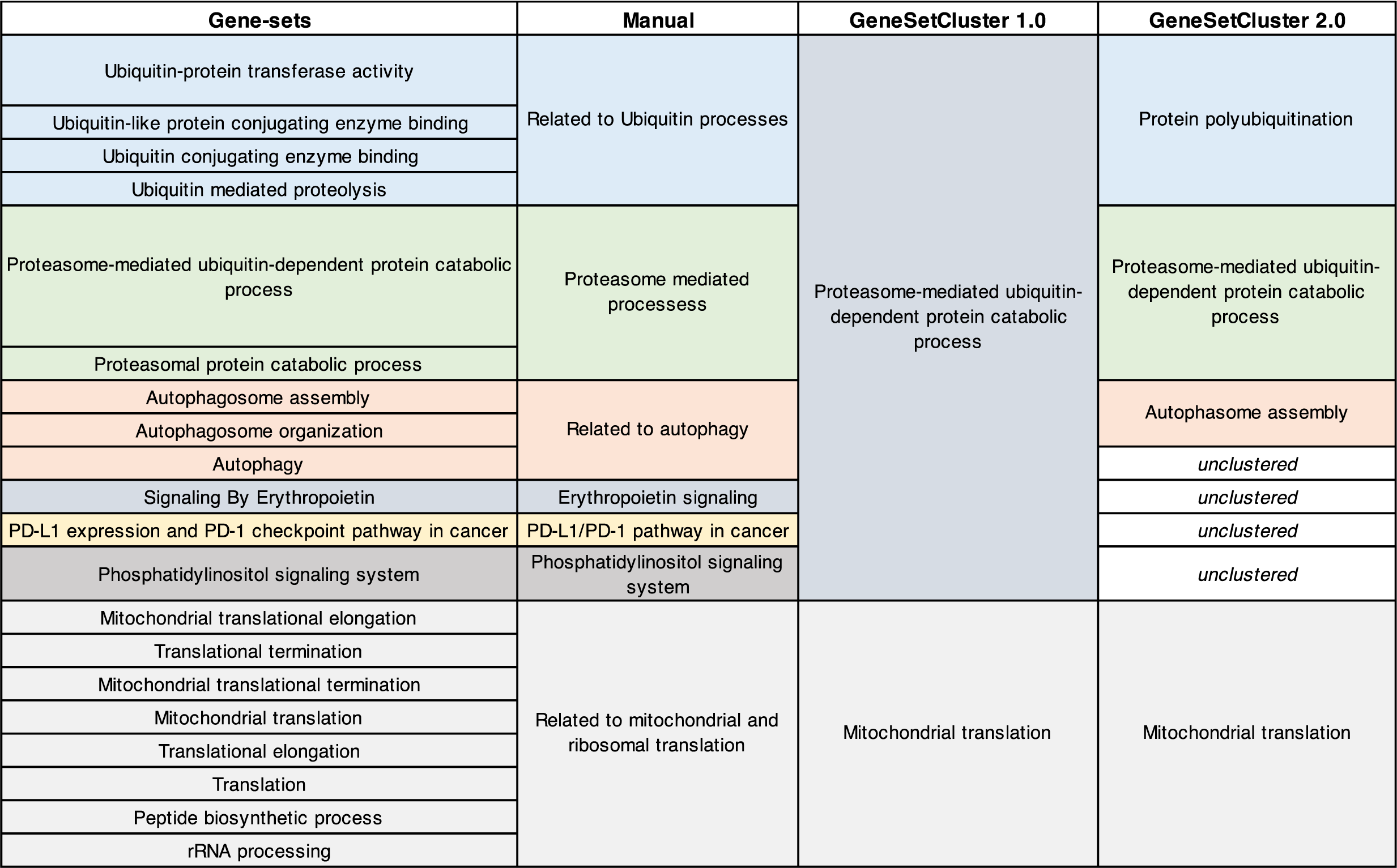
Performance comparison from GeneSetCluster 1.0 and GeneSetCluster 2.0. The first column shows the 20 gene-sets being analyzed. The “Manual” column presents the clusters manually curated by experts. The “GeneSetCluster 1.0” column displays results generated using the “*Raw Gene-sets*” and “*Classic*” methods. The “GeneSetCluster 2.0” column shows the improved results using the “*Unique Gene-sets*” and “*Seriation-based*” methods in GeneSetCluster 2.0.

### 4.2 Reducing computational times

To assess the effectiveness of the parallelization scheme, the *CombineGeneSets* function was tested using datasets of various sizes: small (239 gene-sets), medium (1000 gene-sets), and large (2287 gene-sets), with different numbers of threads: 1, 2, 4, 6, 8, and 10. The execution time was measured for each condition repeated ten times. The tests were conducted on a workstation equipped with Apple M1 Pro processor with a 10-core CPU and 16 GB of RAM.

The mean execution times (in seconds) for each configuration are shown in Table S2. Results showed that performance was slowest with a single thread, but execution times decreased significantly as threads were added, especially for medium and large datasets, up to four threads. For instance, the medium dataset’s processing time dropped from 125 seconds with one thread to 42 seconds with four. Similarly, the large dataset achieved substantial gains, reducing processing time to 440 seconds with six threads, down from 1945 seconds with a single thread. This pattern suggests that using four to six threads effectively balances computational load and efficiency in GeneSetCluster 2.0.

By implementing parallelization, GeneSetCluster 2.0 enhances the scalability and speed of the *CombineGeneSets* step, allowing users to analyze large datasets more effectively.

## 5. Discussion

High-throughput technologies are crucial for profiling biological systems, but interpreting the large number of features identified can be overwhelming. Gene-Set Analysis (GSA) simplifies this by grouping related genes for functional interpretation. Common methods include Over-Representation Analysis and Gene-Set Enrichment Analysis, which assess the importance of gene-sets. However, redundant and overlapping processes often make GSA results difficult to interpret. To overcome this challenge, we developed GeneSetCluster 1.0 (21) using a data-driven approach that clustered gene-sets and enable the biological interpretation of those clusters. Despite its successful application (22,23) GeneSetCluster 1.0 had limitations, including non-interpretable clusters and accessibility barriers. As a result, we developed GeneSetCluster 2.0 to overcome such challenges by improving three areas: methodological development, improved computational times, and a web-based user-friendly version of the tool.

The methodological improvements in GeneSetCluster 2.0 can be directly observed in its ability to avoid the over-aggregation of unrelated gene-sets that occurred in the previous version. The tool more effectively separates gene-sets by combining the “*Unique Gene-sets*” approach and the “*Seriation-based*” clustering method, allowing biologically related processes to be grouped with greater exactness. For example, when GeneSetCluster 1.0 combined 12 gene-sets into a broad *proteasome-mediated processes*, GeneSetCluster 2.0 refined this into biologically distinct clusters, including *protein polyubiquitination*, proteasome-mediated processes, and *autophagosome assembly*. This additional granularity in the clustering enables more precise biological interpretations. The fact that GeneSetCluster 2.0 left three gene-sets unclustered ensures that unrelated gene-sets are not forced into clusters, maintaining a higher degree of biological relevance; furthermore, those non-clustered gene-sets can be investigated separately if required. Thus, v2.0 version demonstrates superior clustering capability, yielding clearer and more meaningful biological insights from complex gene-set data.

Another key improvement is the significant reduction in computational times achieved through optimized parallelization in GeneSetCluster 2.0. The *CombineGeneSets* step, which calculates distances between gene-sets, now leverages parallel processing techniques. By distributing the workload efficiently across multiple processors, the tool achieves faster execution times, especially for larger datasets. This optimization enhances scalability and ensures users can analyze extensive data more effectively.

Finally, incorporating the web-based version makes GeneSetCluster 2.0 more accessible to users with limited bioinformatics expertise. Similar to tools developed in other areas, such accessibility promotes collaboration and more efficient communication within multidisciplinary teams (42–44). Furthermore, a standout feature of GeneSetCluster 2.0 is the seamless integration between the R package and the web application enabling users to transition their analyses and results between platforms effortlessly. For instance, analyses initiated in the R package can be saved and uploaded into the web application to continue exploration with its intuitive interface. Similarly, work started in the web application can be exported back to R for more detailed or customized workflows. This two-way compatibility ensures that users can tailor their workflow to suit their preferences and expertise, facilitating real-time collaboration and efficient sharing of insights.

Despite the novelties and improvements introduced in version 2.0, the discoveries made using GeneSetCluster remain inherently limited to known pathways and biological processes. To develop the framework to overcome such limitations, future developments should focus on integrating exploratory tools that relate gene-set clusters to phenotype information in a more dynamic and exploratory manner, similar to existing tools (45). However, no existing tool provides the precise framework and flexibility that GeneSetCluster 2.0 does.

It is noteworthy that, even after more than 20 years since the inception of GSEA, it is still in development (46). Furthermore, significant advancements are still required to facilitate deeper yet low-complexity biological interpretations. Responding to this need, GeneSetCluster 2.0 improves upon version 1.0 and brings a unique exploratory framework to the field, enabling researchers to characterize gene-sets and their associated clusters better.

## 6. Conclusion

In conclusion, GeneSetCluster 2.0 represents a significant advancement in gene-set analysis interpretation for the research community. Methodologically, it enhances the identification of gene-set clusters and their exploration. Computationally, it ensures faster processing of large datasets. Most importantly, the new web application improves accessibility for researchers with varying levels of coding expertise. Integrating the web application with the R package fosters collaboration between bioinformaticians and biologists, supporting multidisciplinary research efforts. The web application is available at https://translationalbio.shinyapps.io/genesetcluster/. The updated R package, comprehensive documentation, and supporting materials can be downloaded from GitHub at https://github.com/TranslationalBioinformaticsUnit/GeneSetCluster2.0.

