## Supplementary Material for "GeneSetCluster 2.0: a comprehensive toolset for summarizing and integrating gene-sets analysis"

**Table S1.** Seriation methods from the *seriation* R package.

| Algorithm | Method |
|---|---|
| Simulated annealing | **ARSA** |
| Branch-and-bound | **BBURCG** |
| Branch-and-bound | **BBWRCG** |
| TSP solver | **TSP** |
| Optimal leaf ordering | **OLO** |
| | **OLO_single** |
| | **OLO_average** |
| | **OLO_complete** |
| Gruvaeus and Wainer | **GW** |
| | **GW_single** |
| | **GW_average** |
| | **GW_complete** |
| MDS | **MDS** |
| | **MDS_metric** |
| | **MDS_nonmetric** |
| | **MDS_angle** |
| Spectral seriation | **Spectral** |
| | **Spectral_norm** |
| QAP | **QAP_2SUM** |
| | **QAP_LS** |
| | **QAP_BAR** |
| | **QAP_Inertia** |
| Genetic Algorithm | **GA** |
| DendSer | **DendSer** |
| Hierarchical clustering | **HC** |
| | **HC_single** |
| | **HC_verage** |

| | HC_complete |
|---|---|
| Rank-two ellipse seriation | **R2E** |
| Sorting Points Into Neighborhoods | **SPIN_NH** |
| | **SPIN_STS** |
| Visual Assessment of Tendency | **VAT** |

**Table S2.** Average execution time (in seconds) for running the *CombineGeneSets* ten times at different threads (1, 2, 4, 6, 8, and 10). The workstation used for running this experiment has an Apple M1 Pro processor with a 10-core CPU and 16 GB of RAM.

| Dataset | Number Geneset | Thread = 1 | Thread = 2 | Thread = 4 | Thread = 6 | Thread = 8 | Thread = 10 |
|---|---|---|---|---|---|---|---|
| small | 239 | 6.34 | 4.92 | 4.88 | 5.92 | 7.40 | 9.21 |
| medium | 1000 | 125.41 | 72.96 | 42.31 | 33.57 | 30.54 | 32.01 |
| large | 2287 | 1945.20 | 849.50 | 491.70 | 440.20 | 500.41 | 602.44 |

### Performance Scoring Details

The performance scoring of seriation algorithms is based on three weighted criteria:

#### 1. Hamiltonian Path Length (40% weight)

The Hamiltonian path length measures the sum of dissimilarities between consecutive objects in the order:

$$L(D) = \sum_{i=1}^{n-1} d_{i,i+1}$$

Where:

- $D$ is the dissimilarity matrix
- $d_{i,i+1}$ is the dissimilarity between consecutive genesets $i$ and $i+1$
- $n$ is the number of genesets

#### 2. Anti-Robinson Form Criterion (40% weight)

The Anti-Robinson form criterion evaluates how well the dissimilarities increase when moving away from the diagonal:

$$L(D) = \sum_{i=1}^{n-1} (n-i)\, d_{i,i+1}$$

Where:

- $D$ is the dissimilarity matrix
- $d_{i,i+1}$ is the dissimilarity between consecutive genesets $i$ and $i+1$
- $n$ is the number of genesets

#### 3. Total Count of Gene-sets in Clusters (20% weight)

This measures the number of gene-sets successfully assigned to clusters:

$$C(G) = \sum_i |C_i|$$

Where:

- $C(G)$ is the total count of genesets in clusters
- $|C_i|$ is the number of genesets in cluster $i$

**Figure S1**: General schema of GeneSetCluster 2.0.

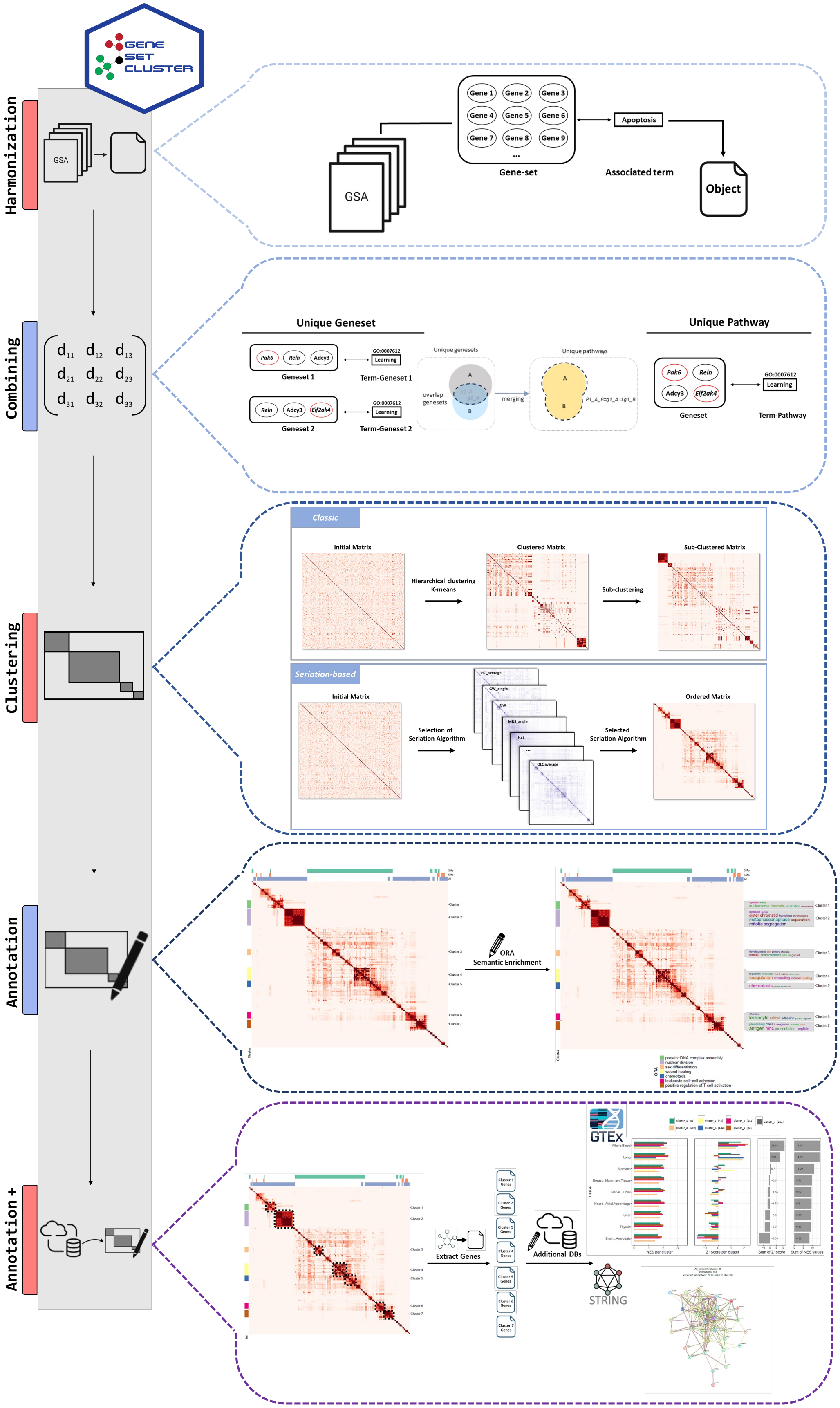
